# Purkinje cell branch morphology determines effect of inhibition and SK2 modulation on somatic pauses

**DOI:** 10.1101/2025.03.31.646261

**Authors:** Pin-Ju Chou, Erik De Schutter, Gabriela Cirtala

## Abstract

Characterized by a highly complex branching of their dendrites, Purkinje cells (PCs) have a unique architecture that enables them to receive impressive amounts of sensorimotor information through their parallel fiber (PF) input. They are tasked to encode this information with high accuracy. In this work, we discuss the mechanisms through which PCs encode this information, and we show how they multiplex between linear-rate and burst-pause coding. Particularly, somatic pauses are of utmost importance due to their involvement in learning. Using a novel heterogeneous model, we show that all branches can achieve a burst-pause response in response to branch-specific PF clustered input. We quantify the somatic pauses obtained and propose various mechanisms to alter the pause duration. Firstly, our results show that increasing local SK2 channel conductance density systematically increases pause duration. In four branches somatic pauses occurred only when SK2 conductance was increased. Interestingly, when adding feed-forward inhibition via stellate cells, our results show either an increase or a decrease in somatic pauses, highlighting the important role of branch morphology and branch location within the PC.

**Significance statement:** Purkinje cells are characterized by highly intricate dendritic branches, which enables them to encode sensorimotor information with great accuracy. Their somatic pauses following excitatory input have been shown to have a strong impact in learning. However, little is known about the impact of morphology and inhibitory input on somatic pauses and implicitly on the learning capacity. In this study, we propose a heterogeneous Purkinje cell model which highlights the importance of branch-specific dendritic morphology on somatic responses. We uncover two different mechanisms for modulating the length of the somatic pauses: density of SK2 channels and feed-forward inhibition via stellate cells.

## 1. Introduction

Purkinje cells (PCs) are one of the largest and most intricate neurons in the brain and are tasked with receiving a wide range of sensory and motor information, generating the only output of the cerebellar cortex. Their unique morphology, with extensive branching of their dendrites, enables PCs to receive more inputs than any other cell type of the brain and allows them to accurately encode precise sensorimotor information. Rat PCs receive approximately 150,000 excitatory synapses from parallel fibers (PFs) (Harvey & Napper, 1991) and their activation triggers local dendritic calcium spikes (Rancz & Häusser, 2006, 2010). Long-term depression (LTD) between PFs and PCs has been intensively studied (Gallimore et al., 2018; Shim et al., 2017) and was shown to be an essential component of motor learning (Ito et al., 1982; Manto et al., 2012; Rancillac & Crépel, 2004). Moreover, studies of LTD based recognition of PF patterns (Steuber et al., 2007) uncovered that the best predictor for pattern recognition learned by LTD was the length of the silent period (pause) after the pattern was presented, showing that the learned patterns resulted in shorter pauses. Over the last decade, somatic pauses and their significant impact in learning were investigated in various experimental studies (Jirenhed et al., 2017; Jirenhed & Hesslow, 2016; Johansson et al., 2016).

Furthermore, studying the mechanisms through which the information transported by spike trains is encoded has been a central topic in neuroscience and has generated a lot of debate over the last few decades (De Zeeuw et al., 2011; Heck et al., 2013; Walter & Khodakhah, 2006). The first evidence of multiplexed coding in cerebellar Purkinje cells was given by Hong et al, who recorded the electrical activity from rhesus monkeys during the execution of fast eye movements (Hong et al., 2016; Markanday et al., 2023). Their results showed that each initiation of eye movement is preceded by a pause in the otherwise regular firing of Purkinje neurons. Therefore, they showed that Purkinje cells use both the timing and the rate of their spiking activity to encode precise movements. Additionally, the role of Purkinje spike burst-pause dynamics has also been examined in a theoretical study on vestibular ocular reflex adaptation (Luque et al., 2019). In this modeling study, we investigate in detail the burst-pause dynamics and we show how the somatic pauses are influenced by biophysical properties.

PC morphological development is tightly linked to its functional maturation, with structural changes directly influencing synaptic integration and circuit refinement (Dusart & Flamant, 2012). During postnatal development, PCs undergo dendritic remodeling and transition from a multiplanar to a monoplanar arrangement (Kaneko et al., 2011; Kato & De Schutter, 2023). This alignment with PF inputs ensures efficient synaptic integration. It is well known that proper dendritic arborization is essential for normal motor behavior and disruptions in this process can lead to motor deficits (Dusart & Flamant, 2012). Moreover, it has been shown that dendritic topology strongly impacts the firing pattern of neurons (de Sousa et al., 2015; Krichmar et al., 2002; Mainen & Sejnowski, 1996; Psarrou et al., 2014; van Elburg & van Ooyen, 2010; van Ooyen et al., 2002). This study showcases the importance of branch-specific morphology on pause duration and, implicitly, on the capacity of processing information.

Besides receiving excitatory input from PFs, PCs also receive crucial inhibitory inputs from molecular layer neurons, primarily basket cells (BCs) and stellate cells (SCs) (Chan-Palay & Palay, 1972; Häusser & Clark, 1997; Heine & Blazquez, 2011; Jelitai et al., 2016; Midtgaard, 1992; Pouzat & Hestrin, 1997; Vincent & Marty, 1996). Inhibitory inputs from BCs and SCs critically sculpt PC firing dynamics (A. M. Brown et al., 2019; He et al., 2015; Hirano & Kawaguchi, 2014; Solinas et al., 2006). Interneurons play a key role in regulating temporal information and shaping the precision of Purkinje cell firing, which is essential for motor coordination (Blazquez & Yakusheva, 2015; Galliano et al., 2013; Jaeger et al., 1996; Wulff et al., 2009) and motor learning (Hirano & Kawaguchi, 2014; Li et al., 2022). Moreover, BCs and SCs serve as the main feed-forward inhibitory pathways in the cerebellum: they receive excitatory input from PFs and, in turn, inhibit PCs at the soma (BCs) and dendrites (SCs), respectively (Bao et al., 2010; Smith & Otis, 2005). Feed-forward inhibition (FFI) is a ubiquitous circuit motif that plays a key role in processes such as sensory perception (Kee et al., 2015; Wang et al., 2018) and motor behavior (Jelitai, 2016). FFI by interneurons increases the temporal precision of spikes (Willadt et al., 2013), filtering out asynchronous excitatory inputs (Mittmann et al., 2005). Stellate cells operate as high-pass filters, inhibiting PCs in response to high-frequency input, thereby regulating their firing and ensuring that only significant signals are transmitted to downstream targets (Mittmann et al., 2005). Moreover, sensory-evoked inhibition has been shown to play a key role in PC processing of external stimuli, with well-timed inhibitory inputs suppressing simple spike activity and modulating cerebellar output (S. T. Brown et al., 2024). Together, these inhibitory interneurons modulate excitatory inputs, ensuring the fine-tuning of PC output necessary for precise motor control (Heine & Blazquez, 2011; Rancillac & Crépel, 2004; Wulff et al., 2009).

We have recently proposed a heterogenous ion channel PC model (Cirtala & De Schutter, 2024) in which each branch of the PC has different ion channel conductance densities, and we have shown how increasing P-type Calcium channel conductance densities (*g*_*CAP*_) facilitates the generation of dendritic calcium spikes (Rancz & Häusser, 2006, 2010), causing a change from linear response to bimodal linear-step-plateau response. We uncovered that each branch has its own characteristic PF threshold, which we defined as the value at which the step response ends. However, this previous work focuses on dendritic responses. In this study, we focus on analyzing the somatic response, and we show how the dendritic calcium spikes evoke somatic pauses that can be modulated by the conductance density of the small-conductance calcium-activated potassium channels (SK2) and via feed-forward stellate cell inhibition. Moreover, we discuss how single branch morphology strongly affects the responses.

## 2. Results

We simulate clustered PF input activation on each of the branches of the heterogeneous PC model (Cirtala & De Schutter, 2024; Zang et al., 2018; Zang & De Schutter, 2021). Background synaptic stimulation of the entire model causes it to spike with a frequency of 24Hz. We show how the heterogeneous PC model (Cirtala & De Schutter, 2024) can multiplex from linear response into burst-pause response at the soma. We quantify the somatic pauses through two different techniques: by analyzing the peristimulus histograms (PSTHs) (Chen et al., 2016) and by determining the inter-spike interval distribution (ISI) (Hong et al., 2016) for activation of each of the branches of the PC model. We apply various statistical techniques to further inspect ISIs and we show how increasing the SK2 conductance density (*g*_*SK*2_) leads to an increase in pause duration. Furthermore, we show the effect that FFI via stellate cells has on the somatic pauses and how morphology impacts this response.

### Somatic response

The first part of the results section presents the somatic responses obtained when simulating clustered PF activation of each branch of the PC using the heterogeneous model proposed by Cirtala and De Schutter (Cirtala & De Schutter, 2024). Particularly, we show how the branch-specific dendritic responses influence the response at the soma.

We illustrate how the heterogenous PC model shifts the linear response into a bimodal response using as an example activation of branch 4 (**Figure 1A-B**). The dendritic responses captured in **Figure 1C-D** show for different numbers of activated PFs, the changes in voltage measured in the distal point p1 for the baseline value of *g*_*CAP*_ (**Figure 1C**) and for a 50% increase in *g*_*CAP*_ that supports dendritic calcium spike generation (**Figure 1D**). The peak amplitude response for the two cases is shown in **Figure 1E** and showcases a linear response for the baseline value of *g*_*CAP*_ (black line) and a bimodal step-plateau response (green line) for the 50% increase in *g*_*CAP*_.

**Figure 1.**
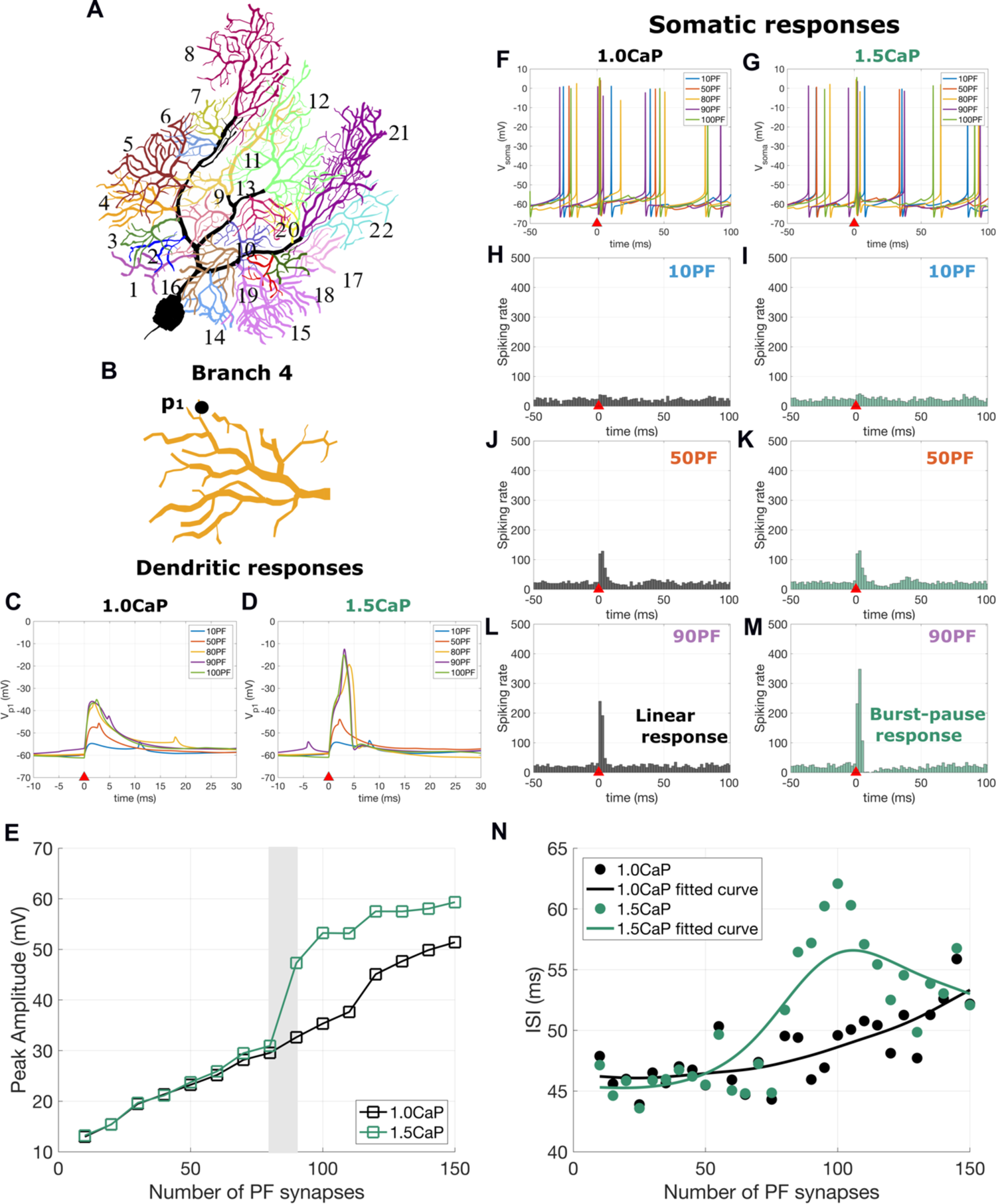
Bimodal linear-step plateau dendritic response results in a burst-pause response at soma. **A.** Morphology of the PC and division into 22 different branches. **B.** Morphology of branch 4. **C and D.** Dendritic calcium spikes recorded at the distal point *p*_1_ for the baseline value of *g*_*CAP*_ (panel C) and for the 50% increase in *g*_*CAP*_ (panel D) for different number of activated PF synapses. In D a shift from EPSP to calcium spike can be observed for activation of 80 or more PFs. **E.** Peak amplitude response for baseline *g*_*CAP*_ (in black) and 50% increase in *g*_*CAP*_ (in green). **F and G.** Voltage measured at soma for baseline and increased *g*_*CAP*_ for activation of 10, 50, 80, 90 and 100 PFs. **H-M**. PSTH responses for 10PF, 50PF and 90PF for baseline (panels H, J and L) and increased *g*_*CAP*_ (I, K and M). **N.** Longest inter-spike intervals (ISI) with increasing number of PF for baseline *g*_*CAP*_ (black) and 50% increase in *g*_*CAP*_ (green). The fitted curves were approximated using a smooth spline.

On the other hand, the right side of **Figure 1** shows the response at soma for these two different scenarios. The voltage recorded at soma is shown in **Figure 1F-G**, while **Figure 1H-M** shows PSTHs for different number of PFs. Observe that for lower number of PFs, the somatic coding response is linear for both *g*_*CAP*_ values (**Figure 1H-K**). Once the threshold value of 90PF is reached, we observe a large burst in the spiking rate, for both baseline and increased value of *g*_*CAP*_ (**Figure 1L** and **Figure 1M**). However, the latter shows a different behavior following the burst: a short pause in the spiking rate (**Figure 1M**). This is known in neural coding as the burst-pause response (Hong et al., 2016; Zang & De Schutter, 2021).

Hence, in this example, we observe that by using the heterogenous ion channel PC model, we were able to shift the dendritic response for branch 4 from linear to bimodal-step-plateau (**Figure 1E**), which resulted in a burst-pause response at the soma for threshold and supra-threshold numbers of activated PF synapses (**Figure 1M**). In **Figure 1N** we show the longest inter-spike interval (ISI) following the PF stimulus with increasing number of PFs for the baseline and increased *g*_*CAP*_. The fitted curves show significant differences in the longest ISIs obtained for the baseline *g*_*CAP*_ compared to the ones obtained for increased *g*_*CAP*_.

We analyzed the somatic responses for each activated branch of the PC model (**Figure 1A**) and we observed that, for most of the dendritic tree, a bimodal dendritic response produces a burst-pause response at soma. **Figure 2A-C** and **Figure 2E-G** show PSTHs of activation of branch 5 and branch 15, in which we can clearly see significant pauses at supra-threshold values. Such behavior occurs in 18 out of the 22 branches (**Figure S2**), with the only exceptions being branch 1, branch 9, branch 10 (**Figure 2I-K**) and branch 22 (**Figure 2M-O**). In these few exceptional cases, we observe that even after the threshold was reached, the somatic burst in spiking rate remains relatively modest and is not followed by a pause.

**Figure 2.**
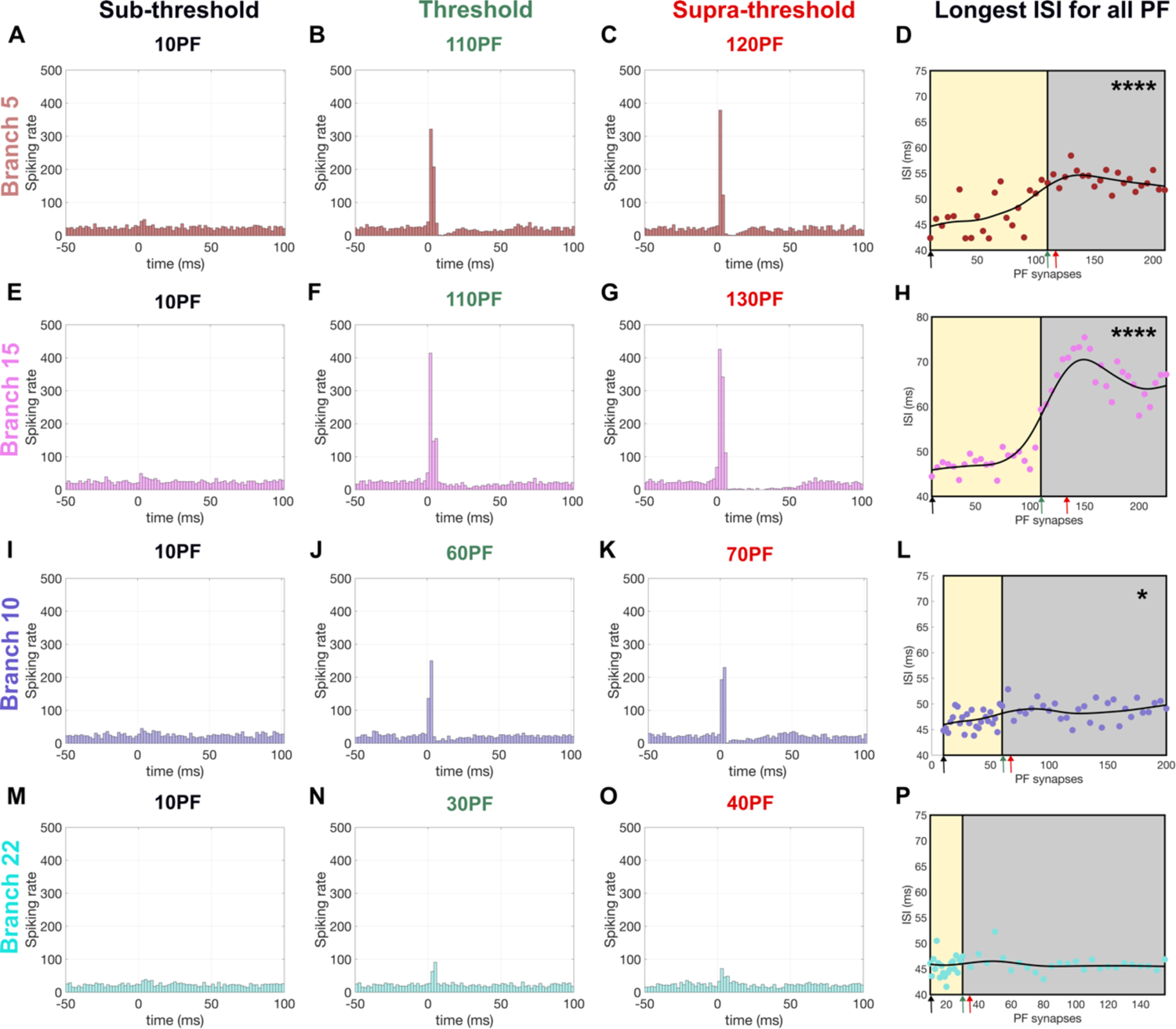
Somatic responses: PSTH and ISIs. In the first three columns we show the PSTHs for three different numbers of activated PF synapses: subthreshold, threshold and supra-threshold for four different branches: branch 5 (**A-C**), branch 15 (**E-G**), branch 10 (**I-K**) and branch 22 (**M-O**). Observe that for all branches, at subthreshold values of PF, we obtain a linear coding response. By increasing the number of PF activated synapses to match the threshold value or beyond, we start observing a burst in the spiking rate, which is in most branches followed by a pause (**B-C** and **F-G**). The last two rows (panels **I-L** and **M-P**) show two exceptions: branches 10 and 22, where no pauses are observed (**J-K** and **N-O**). The last column (panels **D, H, L** and **P**) shows scatter plots of the longest ISIs for the four branches for increasing number of activated PF synapses. The continuous black line represents the fitted data using a smooth spline function. The yellow rectangles contain all PF numbers that are below the threshold value corresponding to each branch, while the gray rectangles comprise of all the supra-threshold PF values. Observe that for the first two branches, as for most of the branches in the dendritic tree, there is a large difference between sub-threshold and supra-threshold, the latter being characterized by much longer maximum ISIs. The three arrows on the bottom of panels **D, H, L** and **P** show the ISI data points corresponding to the PSTH shown on the left side: black arrow (sub-threshold value), green arrow (threshold value) and red arrow (supra-threshold value). The stars on the top right corner of panels **D, H, L** and **P** indicate the p-values obtained for each branch, where we used the following notations: ∗ for p<0.05, ∗∗ for p<0.01, ∗∗∗ for p<0.001 and ∗∗∗∗ for p<0.0001. Note that branch 22 has no star, as its corresponding p-value is larger than 0.05.

To analyze these results in more detail and to visualize these patterns more efficiently over multiple values of activated PF synapses, we devised a strategy different from PSTHs by measuring the ISI duration after the activation of the PF stimulus and picking the longest one (see Methods). We performed 500 trials for each PF value, and we calculated the average of these longest ISIs (Figure S1), which we then show in the scatter plots. The longest ISI scatter plots for the four branches discussed are shown in **Figure 2D** (branch 5), **Figure 2H** (branch 15), **Figure 2L** (branch 10) and **Figure 2P** (branch 22). The ISI data corresponding to each branch is fitted with a smooth spline (black line). The ISI points corresponding to the PSTHs are indicated using black, teal and red arrows on the x-axis and correspond to the sub-threshold, threshold and supra-threshold scenarios shown in the panels on the left side.

To analyze the ISI data, we split the ISIs into two groups: group A shown in the yellow rectangle (**Figure 2** panels D, H, L and P) contains the longest ISI recordings before the branch-specific threshold is reached, and group B with the data shown in the gray rectangle comprises of the longest ISIs after the threshold. We used a two-sided Wilcoxon ranksum test with equal population size, which compares the populations through their medians. Our results capture that, for most of the dendritic tree, the median of the sub-threshold population A is significantly different from the median of supra-threshold population B. However, there are four branches that do not follow the same pattern: branches 1, 9, 10 and 22 show no statistically meaningful difference between the two groups. All statistical information regarding this analysis is given in **Table S1** and the averaged longest ISI data corresponding to every branch within the PC are shown in the Supplementary Material (see **Figure S2**). We believe that the absence of ISI modulation in these 4 branches is due to their very small sizes (see Figure 6 in (Cirtala & De Schutter, 2024)) and the branch position within the tree. In the next sections we show how changing the biophysical properties or/and adding inhibition via stellate cells produces a burst-pause response even in these four branches.

### 2.1. Role of SK2 in somatic pause regulation

According to experimental literature (Kakizawa et al., 2007; Sah, 1996), the activation of the calcium-activated potassium SK2 channel underlies the afterhyperpolarization during action potentials. In addition, the pioneering experimental study by Ohtsuki et al., who used triple patch recordings from different dendritic sites and soma, as well as confocal calcium imaging, shows that the SK2 channel provides a brake mechanism that influences dendritic processing and that such a form of plasticity of dendritic intrinsic excitability can be confined to different activated compartments (Ohtsuki et al., 2012). In this section, we altered the SK2 conductance density, and we determined the effect on both dendritic and somatic responses.

For the four branches that we discussed above, we gradually increased the SK2 conductance density with a parameter α_*SK*2_ such that *g*_*SK*2_ = α_*SK*2_*g*^-^_*SK*2_ where *g*^-^_*SK*2_ represents the reference SK2 conductance density value (Cirtala & De Schutter, 2024; Zang & De Schutter, 2021). In **Figure 3A-D** we visualize the scattered longest ISI plots for branches 1, 9, 10 and 22: with black markers we show the baseline SK2 scenario, while in light blue and red markers we show 2- and 3-fold increases. Observe that in population A, for all branches, the longest ISIs remain relatively unchanged, while population B shows large ISI increases with raising α_*SK*2_.

**Figure 3.**
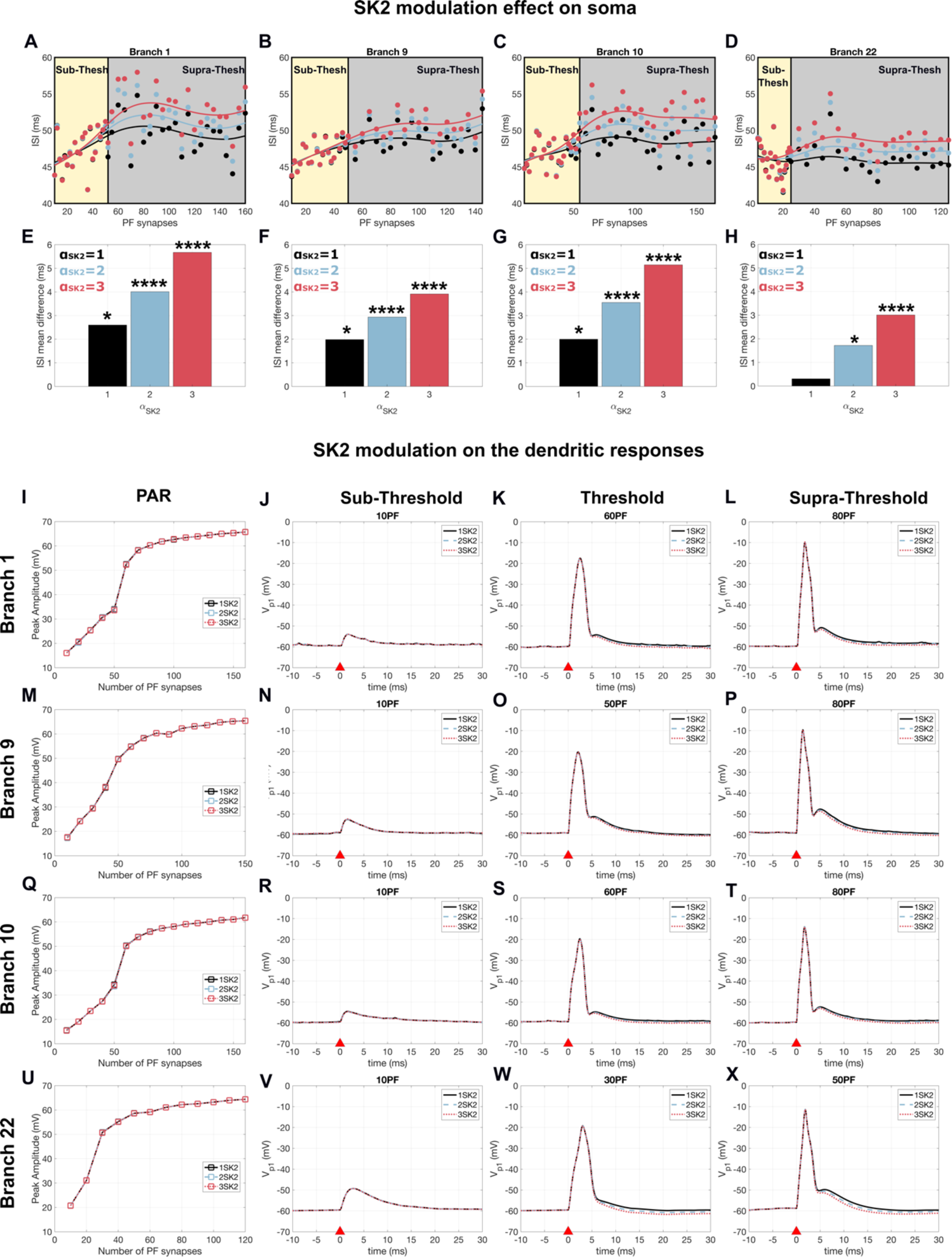
SK2 modulation effect on somatic and dendritic responses. In panels **A-D** we show the scatter plots of the longest ISIs recorded for branch 1, 9, 10 and 22. They are fitted with a smooth spline function. The yellow rectangle shows the averaged longest ISIs over all the trials belonging to population A, which contains the ISI values before the threshold is reached while the gray rectangle captures population B, the ISI values obtained after the threshold is reached. The bar plots in panels **E-H** show the average longest ISI difference for different increases in α_*SK*2_. The stars on the bar plots in panels **E-H** show the p-values obtained for each branch, where we used the following notations: ∗ for p<0.05, ∗∗ for p<0.01, ∗∗∗ for p<0.001 and ∗∗∗∗ for p<0.0001. The second part of this figure focuses on the dendritic responses. Showing the peak amplitude response and voltage traces for sub-threshold, threshold and supra-threshold numbers of PF for different increases in α_*SK*2_ for branches 1 (panels **I-L**), 9 (panels **M-P**), 10 (panels **Q-T**) and 22 (panels **U-X**).

When comparing the average longest ISI difference between the sub-threshold and supra-threshold populations, we observe substantial differences for all four branches (**Figure 3E-H**). The statistical analysis (see **Table S2**) shows that increasing the conductance density of SK2 channels produces a significant increase in the mean of population B corresponding to a pause in firing, while population A remains relatively constant. Therefore, when we examined the difference between these two populations, we observed that increases in α_*SK*2_ lead to meaningful statistical differences (p-value < 0.01) for all four branches, therefore showing that they exhibit a burst-pause response. More details regarding the statistical analysis can be found in Methods and all statistical results are presented in **Table S2**.

In the second part of **Figure 3**, we show the effect that increased α_*SK*2_ has on the dendritic responses of each of the four analyzed branches. Note that the peak amplitude response remains unchanged (**Figure 3** panels **I, M, Q** and **U**), therefore the dendritic bimodal-step plateau response is maintained for different α_*SK*2_. The voltage traces for low number of activated PF synapses remain unchanged for different values of α_*SK*2_ (**Figure 3** panels **J, N, R** and **V**). However, when increasing the PF number to threshold and supra-threshold values, we observe a modest increase of the after-hyperpolarization phase with increasing α_*SK*2_.

Therefore, our results agree with the experimental work by Grasselli et al., according to which the SK channel downregulation reduces AHP amplitude, as well as the pause duration (Grasselli et al., 2016).

### 2.2. Feed-forward inhibition by stellate cells

In this section, we describe the effects of adding dendritic FFI to the somatic and dendritic responses. We included FFI to the heterogeneous model by adding stellate cells and we activated each branch of the dendritic tree. The inhibitory input was added 1.4 ms after the activation of the excitatory PF synapses, which represents the average EPSP-IPSP delay for FFI recorded in experiments (Mittmann et al., 2005).

#### 2.2.1. Inhibition shifts the PF threshold

First, we examined the response to FFI when activating 10 to 150 PF synapses and 0, 75 or 150 stellate cell synapses. The peak amplitude response obtained for all branches is shown in Figure 4. The black lines and markers correspond to the baseline case in which no inhibition was added, while the cyan and purple ones correspond to 75 and 150 stellate cell synapses, respectively. We observed that most branches exhibit a shift of the PAR curve to the right side with increasing inhibition. Such behavior was also reported in Zang et al. when using a homogeneous model (Zang & De Schutter, 2021). In Figure 4 we marked the “jump” response of each branch using light gray rectangles, while the dark gray rectangles show the rightwards shifting effect.

**Figure 4.**
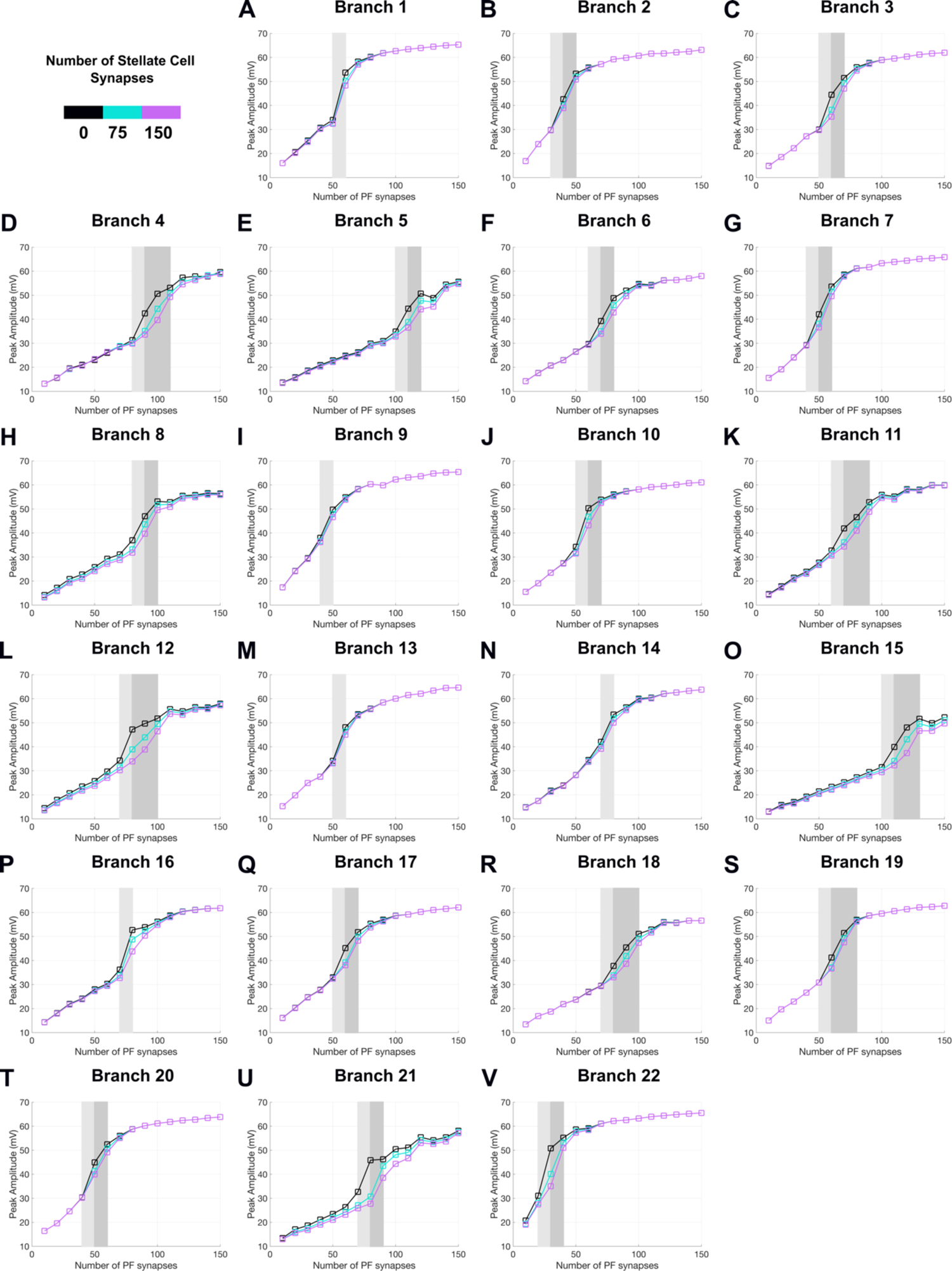
Feed-forward Inhibition by stellate cells shifts the PF threshold. We show the peak amplitude response for each of the 22 branches for three different scenarios: no inhibition (black), 75 stellate cells synapses (cyan) and 150 stellate cell synapses (purple). The light gray rectangle captures the jump response without inhibition. The threshold characteristic to each branch is defined as the PF number where the jump ends. The dark gray rectangle shows the rightwards branch-specific shifting effect of inhibition.

For example, with no inhibition, branch 4 (Figure 4D) has a linear response until 80PF, after which it exhibits a jump caused by dendritic calcium spikes at a threshold of 90PF. When adding FFI by 75 stellate cells synapses, the peak amplitude response slightly shifts rightwards, the linear response now ends at 90PF, and it jumps at a threshold of 100PF (cyan line in Figure 4D). When increasing inhibition further, the shift is even more pronounced, with the “jump” occurring at 110PF.

This shifting effect, however, is not prevalent in all branches: for example, no shift could be detected in branch 1 (Figure 4A), 9 (Figure 4I), 13 (Figure 4M) or 14 (Figure 4N). Moreover, each branch has its own unique increment in dendritic spike threshold. These thresholds are very crucial in the soma analysis, as we will explain in the upcoming subsection 2.2.2.

#### 2.2.2. Inhibition effect on somatic pauses

To investigate the effect of inhibition at the soma, we employed the same strategy as in subsection 2.2.1. We separated the ISI data into two groups, and we performed statistical analysis by using the two-sided Wilcoxon ranksum test as described in the Methods section. Due to the shifting effect shown in the previous subsection, showcasing large changes in PF threshold for different levels of inhibition, we excluded the shift-zone (gray rectangles in Figure 4) from the statistical analysis.

Our simulations show that adding FFI via stellate cells systematically increase ISIs for both populations studied (sub-threshold Figure 5C-D and supra-threshold Figure 5A-B). However, when computing the difference between population B and population A, we observed that while most of the tree shows clear increases in ISIs, there are a few branches on the right side of the PC (Figure 5E) for which inhibition acts conversely, decreasing ISIs (Figure 5F-G). Possible explanations of this masking phenomenon are provided in the Discussion.

**Figure 5.**
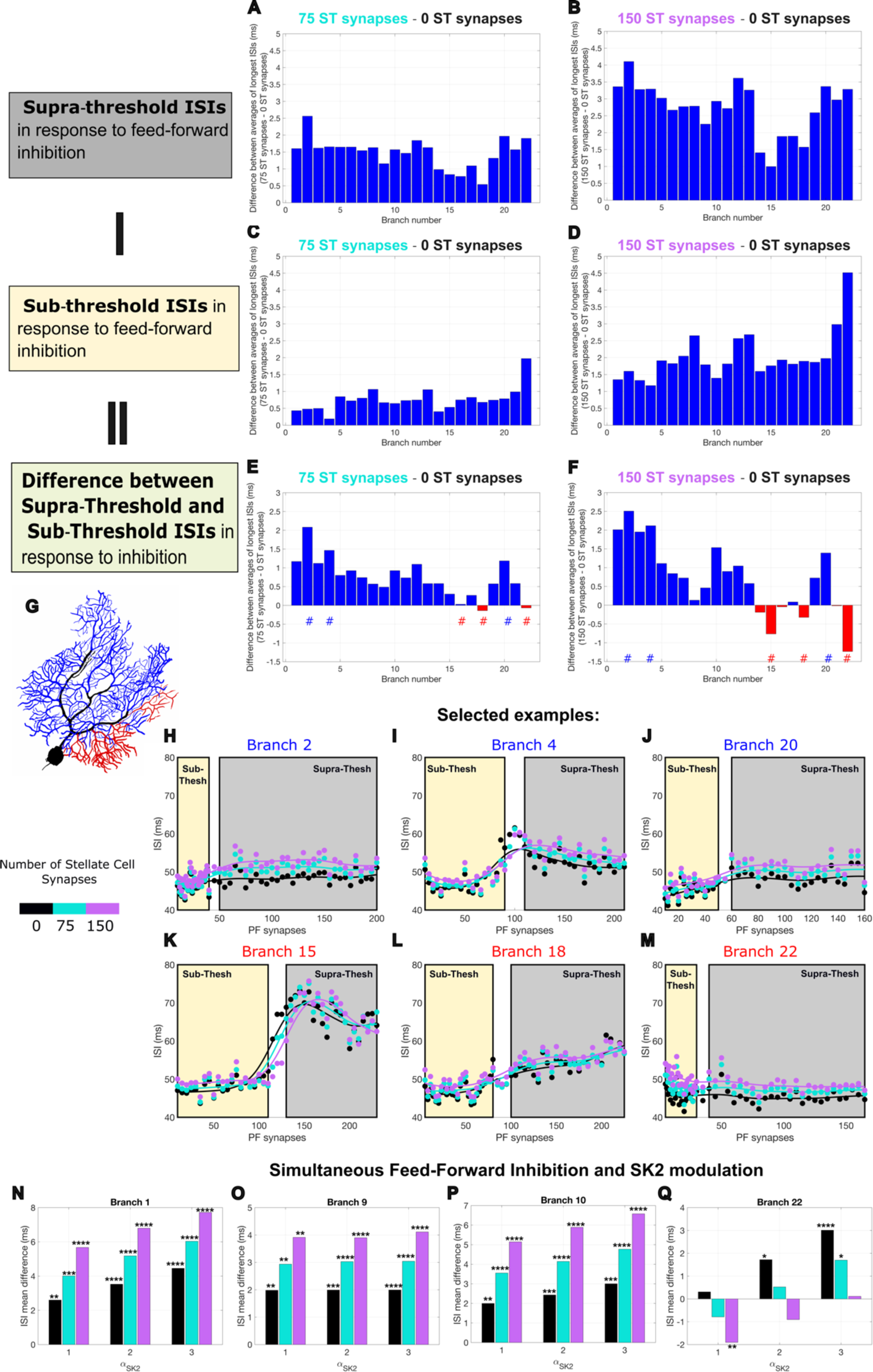
The impact of feed-forward inhibition on somatic pauses. **A-B.** Longest ISIs calculated for supra-threshold values for all branches: panel A shows the ISI difference between 75 stellate cell synapses and no inhibition scenario, while panel B captures the ISI difference between 150 stellate cell synapses and no inhibition. **C-D.** Longest ISI differences for sub-threshold PF values. **F-G.** Computed difference between the cases shown above: panel E represents the computed ISI difference between panel A and panel C, while panel F shows the ISI difference between panel B and D. The results show that for most branches, FFI results in an increase in pause duration (blue), though for a few exceptions, FFI can also decrease pauses (red). The location of these two branch categories can be visualized in panel **E**. Observe that all branches whose FFI decreases pauses (red) belong to the right division of the PC. **H-M.** Examples of branches that exhibit the highest increases in ISI (branches 2, 4 and 20) when adding FFI (panel H-J) and branches that show large decreases (branches 15, 18 and 22 in panels K-M). **N-Q.** Longest ISI mean difference for simultaneous FFI and SK2 modulation. The amount of inhibition is shown in colors: black (no inhibition), cyan (75 stellate cell synapses) and purple (150 stellate cell synapses), while the SK2 increases are shown on the x-axis. The stars on top of the bar plots in panels **N-Q** indicate the p-values obtained for each branch, where we used the following notations: ∗ for p<0.05, ∗∗ for p<0.01, ∗∗∗ for p<0.001 and ∗∗∗∗ for p<0.0001.

In Figure 5H-M we show the longest ISI data scatter plots for a few branches that had the highest increases and decreases in the ISI difference. For example, branch 2 had the highest ISI increase for the maximum FFI considered (150 SC synapses). Observe in the yellow box the small ISIs obtained sub-threshold and note when we increase the amount of inhibition, the difference between the sub-threshold and supra-threshold becomes much larger (Figure 5H-J). On the other hand, the largest decrease in ISI can be seen in branch 22. Indeed, in Figure 5M, we can visualize that the ISIs in the sub-threshold group (yellow rectangle) are larger than the ones in the supra-threshold one (gray rectangle). All statistical information for different levels of inhibition can be found in **Table S3**.

### 2.3. SK2 and feed-forward inhibition

In this subsection we present the results obtained when modulating α_*SK*2_ for different levels of inhibition. In the colored bar plots in Figure 5N-Q we observe the mean ISI difference for baseline *g*_*SK*2_, for a 2-fold increase and a 3-fold increase. For all four branches analyzed, we observe that increasing SK2 systematically increases ISIs for all branches. As we increase inhibition simultaneously with SK2, the longest ISIs increase for 3 of the branches studied (branches 1, 9 and 10). However, for branch 22, as observed in the previous section in the case of adding inhibition, we observe a decrease in ISI as we increase the number of stellate cells considered.

## 3. Discussion

Our results show that PCs employ branch-specific multiplexed coding. We revealed how a dendritic bimodal step-plateau response translates to a burst-pause response at the soma when using the heterogeneous ion channel model with increased conductance density of CaP and SK2. Without this heterogeneity, we have shown that a linear dendritic response translates to a linear response at soma. PCs use such multiplexed coding to increase their cerebellar coding and learning capacity, which is in agreement with previous experimental and computational work (Hong et al., 2016; Zang & De Schutter, 2021).

In section 2.2 we show that increasing α_*SK*2_ systematically increases ISIs (Figure 3) and therefore pause duration. Similar behavior was observed experimentally by Grasseli et al. when studying complex spikes pauses (Grasselli et al., 2016). To our knowledge, this is the first study that includes SK2 heterogeneity in a computational model. There is however extensive experimental evidence of SK2 heterogeneity in Purkinje cells (Grasselli et al., 2016; Ohtsuki, 2020; Ohtsuki et al., 2012, 2020; Ohtsuki & Hansel, 2018). In particular, a recent article (Ohtsuki et al., 2020) emphasizes the importance of SK mediated non-synaptic form of intrinsic plasticity and shows how these mechanisms can enhance the computational power of neurons through branch specific excitability.

When adding feed-forward inhibition, we have seen that for both sub-threshold and supra-threshold ISI data, adding inhibition via stellate cells increases pause duration. However, when comparing the difference between these two populations, we observed that, while for most branches inhibition increases pause duration, there are a few exceptions for which adding larger levels of inhibition can provoke a slight decrease in ISI, masking the effect of the dendritic calcium spike. These few branches were entirely localized on the right-most division of the PC. We hypothesize that branch morphology, branch location within the tree and the degree to which the dendritic calcium spikes remain constrained within the activated branch play an important factor with respect to ISI length. In fact, our previous work (Cirtala & De Schutter, 2024) shows that most dendritic calcium spikes are relatively well constrained, except for the right main division of the PC, where dendritic calcium spikes tend to spread towards the nearby branches. We have also shown that this might be due to fact that the lengths between different branches are much smaller in this side of the tree (Figure S2 in (Cirtala & De Schutter, 2024)).

Therefore, this study showcases the importance of branch-specific morphology on inhibition and on cerebellar coding. The significant role that morphology plays on neuronal firing has been uncovered in different neuron types over the last decades (de Sousa et al., 2015; Krichmar et al., 2002; Mainen & Sejnowski, 1996; Psarrou et al., 2014; van Elburg & van Ooyen, 2010; van Ooyen et al., 2002). However, the relationship between branch-specific morphology and inhibition has not yet been studied, to our knowledge, in any theoretical or experimental setting.

## 4. Conclusions

Starting from the branch-specific heterogeneous model (Cirtala & De Schutter, 2024), we have analyzed the somatic responses in detail, and we have shown how the bimodal step-plateau response recorded at the tip of each dendritic branch translates into a burst-pause somatic response. While in previous work (Zang & De Schutter, 2021), only 4 branches were achieving this burst-pause somatic response, this novel heterogeneous model shows that all branches can multiplex when adding SK2 heterogeneity. In addition, we have shown that FFI via stellate cells has a strong impact on the somatic pause duration, increasing it for most branches, while decreasing it for few exceptions in the right side of the tree. Our results showcase the importance of branch-specific morphology on the somatic response and therefore, suggest a large impact on learning.

## 5. Limitations of Study

When simulating FFI via stellate cells, we considered 3 different levels of inhibition: 0 ST, 5 ST and 10 ST, with each ST forming 15 synapses on the spiny dendrites. We considered a maximum of 10 ST due to experimental rat studies (Korbo et al., 1993) showing that the ratio between ST to PC is approximately 10:1. We limited ourselves to choosing only three values of inhibition due to the very large number of simulations involved: for each of the 22 branches, three distinct levels of inhibition for the large range of PF synapses explored yielded approximately 75,000 simulations. Additional inhibition levels could in theory bring more information on how FFI affects pause duration, particularly for the branches in the right side of the tree, for which adding the maximum 10 ST inhibition resulted in a decrease of its somatic pause.

Additionally, our inhibition protocol is limited to stellate cells and does not include basket cells. Contrary to stellate cells, which form synapses solely on the PC dendritic tree, basket cells surround the cell bodies of PCs and, therefore, are not involved in the generation of dendritic spikes. However, inhibition by basket cells will strongly contribute to somatic pauses.

Moreover, in this study we did not include any variation in the somatic current injection, our PC corresponding to the zebrin positive module. We have previously shown for climbing fiber excitatory input that PCs located in the Zebrin negative module, characterized by much higher frequencies of 76 ± 19.5Hz (Zhou et al., 2014), exhibit a much shorter pause than their counterparts in zebrin positive (Cirtala & De Schutter, 2024; Zhou et al., 2014). However, the experimental work by Zhou et al shows no significant changes in the waveform and simple spike regularity in different zebrin zones (Zhou et al., 2014).

## 6. Methods

The multi-compartment model we propose is based on a recent heterogenous model (Cirtala & De Schutter, 2024) and comprises of soma, axon initial segment, and smooth and spiny dendrites. The detailed morphology used belongs to a 21 days old Wistar rat PC (Roth & Häusser, 2001). Each of the four sections is characterized by different ion channel conductance densities. In addition, each branch of the PC is characterized by a different P-type calcium channel conductance density, as well as Kv4 channel conductance density as specified in (Cirtala & De Schutter, 2024). In addition, we have also included heterogeneity in SK2 channel conductance density for some branches, as discussed in the Results section. Moreover, we simulated FFI via ST, with each ST forming 15 synapses on the spiny dendrites of the PC. For the inhibitory input, we implemented a 1.4ms delay after the PF activation to account for the average EPSP-IPSP delay for FFI recorded in experiments. All simulations were implemented in NEURON 8.2. For the data analysis, statistics and plots we used MATLAB R2022a and Python 3.7. The NEURON code and the data analysis code are publicly available on ModelDB.

### General simulation protocol

To investigate the somatic responses, we used the same simulation protocol as in (Zang & De Schutter, 2021): we simulated 500 trials for each distinct number of PFs = [10,200]. Each trial consists of background excitatory and inhibitory synapses randomly activated and the somatic timing was disturbed by a holding current of different lengths. For a complete description of this approach and how adding background synaptic inputs leads to very accurate dendritic membrane potential variations when compared to in vivo recordings, please see (Zang & De Schutter, 2021).

### Statistical analysis methods

We analyzed the somatic responses using two different approaches: via PSTH, as done in (Zang & De Schutter, 2021) and via ISIs, as proposed in (Hong et al., 2016). For plotting PSTHs we used a bin size of 2 milliseconds and the data from the 500 trials described above. The main disadvantage of the PSTH technique is the averaging component. To increase precision and to observe the full range of simulated PF synapses we computed the longest ISIs, and we used statistical techniques to examine the differences between different populations.

We measured the intervals between different spikes at soma after the activation time of the PF synapses. For each PF and for each trial, we calculated all ISIs obtained within a time frame of 120 ms post-activation and we selected the longest one (see example in **Figure S1A**). This resulted in 500 different ISIs for each PF (**Figure S1B**). We calculated the average over all trials and we obtained one longest ISI for each PF. We repeated this procedure for all branches for PF in [10:200]. We analyzed the longest ISIs using the two-sided Wilcoxon ranksum test, which compares two different populations through their medians. The two populations we considered included a sub-threshold population which contains the averaged longest ISIs obtained for PF values smaller than the branch-specific characteristic threshold and a supra-threshold population. The threshold value was excluded from our analysis. For accuracy, we used equal population sizes and a minimum data set of 20 samples. Note that each branch has its own characteristic PF threshold (see Figure 4, light gray rectangle), and therefore we varied the PF step, and the maximum PF number considered such that size of the sub-threshold ISI population and the size of supra-threshold ISI population is equal and such that each population has a minimum 20 ISI data points. We used a confidence level of 99% and therefore rejected the hypothesis for any p-value larger than 0.01.

The p-values corresponding to each branch for each case examined are indicated in the figures using the star notation: ∗ for p<0.05, ∗∗ for p<0.01, ∗∗∗ for p<0.001 and ∗∗∗∗ for p<0.0001. The panels where no star is present denote a p≥0.05. The corresponding p-values for each case are presented in detail in the Supplementary material.

### Computing ISIs during different protocols

We have seen in section 2.2 that increasing the conductance density of SK2 does not change the bimodal response and therefore, within the same branch, the PF threshold does not vary with increasing SK2. Therefore, we could employ the same ISI computing protocol as described above. However, in section 2.3, we observed that adding FFI alters the bimodal response, and consequently the PF threshold, for most branches. Hence, the ISIs corresponding to the shifting period marked in dark gray rectangles in Figure 4 were excluded from our analysis. For example, branch 4 (Figure 4D) has a PF threshold of 90PF for no inhibition, a threshold of 100PF when inhibited via 75 ST synapses and a threshold of 110PF when considering 150 ST synapses. Therefore, the sub-threshold population is characterized by all ISIs obtained for PF numbers larger or equal than 10 and smaller than 90, while the supra-threshold population comprises ISIs for PFs in the range [110,210].

### SK2 simulations

For the branches that showed no meaningful statistical difference between the two populations, we increased α_*SK*2_ by 2 and 3-fold. The ISIs corresponding to different SK2 values are shown in Figure 3 and are fitted using a smooth spline with s = 0.0001. In the dendritic responses shown in Figure 3I-X for different values, we averaged 20 randomly selected trials out of the 500 simulations. All statistical information is given in **Table S2**.

### Feed-forward-inhibition (FFI)

The simulation protocol regarding FFI consists of adding 0, 5 and 10 ST, each comprising of 15 synapses onto the activated branch. These different inhibition inputs were added 1.4 ms after the activation of excitatory PF synapses as done in experimental work (Mittmann et al., 2005). Figure 4 shows the effect of inhibition on dendritic responses when averaging 20 randomly selected trials.

## Acknowledgements

This work is supported by the Okinawa Institute of Science and Technology Graduate University. We thank the Scientific Computing & Data Analysis Section (SCDA) of OIST for providing access to the OIST computing cluster.

## Author Contributions

GC, EDS and PC conceived this study. GC and PC performed all simulations. PC, EDS and GC analyzed the data. PC, EDS and GC wrote the manuscript.

## Declaration of interests

The authors declare no competing interests.

## Notes

### Competing Interest Statement

The authors have declared no competing interest.

## References

Bao, J., Reim, K., & Sakaba, T. (2010). Target-dependent feedforward inhibition mediated by short-term synaptic plasticity in the cerebellum. Journal of Neuroscience, 30(24), 8171–8179. 10.1523/JNEUROSCI.0276-10.2010

Blazquez, P. M., & Yakusheva, T. A. (2015). GABA-A Inhibition Shapes the Spatial and Temporal Response Properties of Purkinje Cells in the Macaque Cerebellum. Cell Reports, 11(7), 1043–1053. 10.1016/j.celrep.2015.04.020

Brown, A. M., Arancillo, M., Lin, T., Catt, D. R., Zhou, J., Lackey, E. P., Stay, T. L., Zuo, Z., White, J. J., & Sillitoe, R. V. (2019). Molecular layer interneurons shape the spike activity of cerebellar Purkinje cells. Scientific Reports, *9*(1). 10.1038/s41598-018-38264-1

Brown, S. T., Medina-Pizarro, M., Holla, M., Vaaga, C. E., & Raman, I. M. (2024). Simple spike patterns and synaptic mechanisms encoding sensory and motor signals in Purkinje cells and the cerebellar nuclei. Neuron, 112(11), 1848–1861.e4. 10.1016/j.neuron.2024.02.014

Chan-Palay, V., & Palay, S. L. (1972). The Stellate Cells of the Rat’s Cerebellar Cortex*. Z. Anat. Entwickl.-Gesch, 13C, 224–248.

Chen, S., Augustine, G. J., & Chadderton, P. (2016). The cerebellum linearly encodes whisker position during voluntary movement. ELife, 5(e10509). 10.7554/eLife.10509.001

Cirtala, G., & De Schutter, E. (2024). Branch-specific clustered parallel fiber input controls dendritic computation in Purkinje cells. IScience, 27(9), 110756. 10.1016/j.isci.2024.110756

de Sousa, G., Maex, R., Adams, R., Davey, N., & Steuber, V. (2015). Dendritic morphology predicts pattern recognition performance in multi-compartmental model neurons with and without active conductances. Journal of Computational Neuroscience, 38(2), 221–234. 10.1007/s10827-014-0537-1

De Zeeuw, C. I., Hoebeek, F. E., Bosman, L. W. J., Schonewille, M., Witter, L., & Koekkoek, S. K. (2011). Spatiotemporal firing patterns in the cerebellum. In Nature Reviews Neuroscience (Vol. 12, Issue 6, pp. 327–344). 10.1038/nrn3011

Dusart, I., & Flamant, F. (2012). Profound morphological and functional changes of rodent Purkinje cells between the first and the second postnatal weeks: A metamorphosis? In Frontiers in Neuroanatomy (Issue MARCH, pp. 1–26). 10.3389/fnana.2012.00011

Galliano, E., Gao, Z., Schonewille, M., Todorov, B., Simons, E., Pop, A. S., D’Angelo, E., Van Den Maagdenberg, A. M. J. M., Hoebeek, F. E., & De Zeeuw, C. I. (2013). Silencing the Majority of Cerebellar Granule Cells Uncovers Their Essential Role in Motor Learning and Consolidation. Cell Reports, 3(4), 1239–1251. 10.1016/j.celrep.2013.03.023

Gallimore, A. R., Kim, T., Tanaka-Yamamoto, K., & De Schutter, E. (2018). Switching On Depression and Potentiation in the Cerebellum. Cell Reports, 22(3), 722–733. 10.1016/j.celrep.2017.12.084

Grasselli, G., He, Q., Wan, V., Adelman, J. P., Ohtsuki, G., & Hansel, C. (2016). Activity-Dependent Plasticity of Spike Pauses in Cerebellar Purkinje Cells. Cell Reports, 14(11), 2546–2553. 10.1016/j.celrep.2016.02.054

Harvey, R. J., & Napper, R. M. A. (1991). Quantitative studies on the mammalian cerebellum. Progress in Neurobiology, *36*, 437–463.

Häusser, M., & Clark, B. A. (1997). Tonic Synaptic Inhibition Modulates Neuronal Output Pattern and Spatiotemporal Synaptic Integration. Neuron, *19*(3), 665–678. 10.1016/S0896-6273(00)80379-7

He, Q., Duguid, I., Clark, B., Panzanelli, P., Patel, B., Thomas, P., Fritschy, J. M., & Smart, T. G. (2015). Interneuron- and GABAA receptor-specific inhibitory synaptic plasticity in cerebellar Purkinje cells. Nature Communications, C. 10.1038/ncomms8364

Heck, D. H., De Zeeuw, C. I., Jaeger, D., Khodakhah, K., & Person, A. L. (2013). The neuronal code(s) of the cerebellum. Journal of Neuroscience, 33(45), 17603–17609. 10.1523/JNEUROSCI.2759-13.2013

Heine, S. A., & Blazquez, P. M. (2011). Thinking Inside the Box: The Roles of Inhibitory Interneurons in Cerebellar Cortical Processing. Journal of the Japanese Society for Neural Networks, 18, 50–58.

Hirano, T., & Kawaguchi, S. Y. (2014). Regulation and functional roles of rebound potentiation at cerebellar stellate cell-Purkinje cell synapses. Frontiers in Cellular Neuroscience, 8(FEB). 10.3389/fncel.2014.00042

Hong, S., Negrello, M., Junker, M., Smilgin, A., Thier, P., & De Schutter, E. (2016). Multiplexed coding by cerebellar Purkinje neurons. ELife, 5, e13810. 10.7554/eLife.13810.001

Ito, M., Sakurai, M., & Tongroach, P. (1982). Climbing fibre induced depression of both mossy fribre responsiveness and glutamate sensitivity of cerebellar Purkinje cells. Journal of Physiology, 324, 113–134.

Jaeger, D., De Schutter, E., & Bower, J. M. (1996). The Role of Synaptic and Voltage-Gated Currents in the Control of Purkinje Cell Spiking: A Modeling Study. In The Journal of Neuroscience (Vol. 19, Issue 1).

Jelitai, M., Puggioni, P., Ishikawa, T., Rinaldi, A., & Duguid, I. (2016). Dendritic excitation-inhibition balance shapes cerebellar output during motor behaviour. Nature Communications, 7. 10.1038/ncomms13722

Jirenhed, D. A., & Hesslow, G. (2016). Are Purkinje Cell Pauses Drivers of Classically Conditioned Blink Responses? In Cerebellum (Vol. 15, Issue 4, pp. 526–534). Springer New York LLC. 10.1007/s12311-015-0722-4

Jirenhed, D. A., Rasmussen, A., Johansson, F., & Hesslow, G. (2017). Learned response sequences in cerebellar Purkinje cells. Proceedings of the National Academy of Sciences of the United States of America, 114(23), 6127–6132. 10.1073/pnas.1621132114

Johansson, F., Hesslow, G., & Medina, J. F. (2016). Mechanisms for motor timing in the cerebellar cortex. Current Opinion in Behavioral Sciences, 8, 53–59. 10.1016/j.cobeha.2016.01.013

Kakizawa, S., Kishimoto, Y., Hashimoto, K., Miyazaki, T., Furutani, K., Shimizu, H., Fukaya, M., Nishi, M., Sakagami, H., Ikeda, A., Kondo, H., Kano, M., Watanabe, M., Iino, M., & Takeshima, H. (2007). Junctophilin-mediated channel crosstalk essential for cerebellar synaptic plasticity. EMBO Journal, *26*(7), 1924–1933. 10.1038/sj.emboj.7601639

Kaneko, M., Yamaguchi, K., Eiraku, M., Sato, M., Takata, N., Kiyohara, Y., Mishina, M., Hirase, H., Hashikawa, T., & Kengaku, M. (2011). Remodeling of monoplanar purkinje cell dendrites during cerebellar circuit formation. *PLoS ONE*, 6(5). 10.1371/journal.pone.0020108

Kato, M., & De Schutter, E. (2023). Models of Purkinje cell dendritic tree selection during early cerebellar development. PLOS Computational Biology, *19*(7), e1011320. 10.1371/journal.pcbi.1011320

Kee, T., Sanda, P., Gupta, N., Stopfer, M., & Bazhenov, M. (2015). Feed-Forward versus Feedback Inhibition in a Basic Olfactory Circuit. PLoS Computational Biology, 11(10). 10.1371/journal.pcbi.1004531

Korbo, L., Andersen, B. B., Ladefoged, O., & Møller, A. (1993). Total numbers of various cell types in rat cerebellar cortex estimated using an unbiased stereological method. Brain Research, 609(1–2), 262–268. 10.1016/0006-8993(93)90881-M

Krichmar, J. L., Nasuto, S. J., Scorcioni, R., Washington, S. D., & Ascoli, G. A. (2002). Effects of dendritic morphology on CA3 pyramidal cell electrophysiology: a simulation study. *Brain Research*, 941(1–2), 11–28. 10.1016/S0006-8993(02)02488-5

Li, W., Chen, L., Fleming, J. T., Brignola, E., Zavalin, K., Lagrange, A., Rex, T., Heiney, S. A., Wojaczynski, G. J., Medina, J. F., & Chiang, C. (2022). Dendritic Inhibition by Shh Signaling-Dependent Stellate Cell Pool Is Critical for Motor Learning. Journal of Neuroscience, 42(26), 5130–5143. 10.1523/JNEUROSCI.2073-21.2022

Luque, N. R., Naveros, F., Carrillo, R. R., Ros, E., & Arleo, A. (2019). Spike burst-pause dynamics of Purkinje cells regulate sensorimotor adaptation. PLoS Computational Biology, 15(3). 10.1371/journal.pcbi.1006298

Mainen, Z. F., & Sejnowski, T. J. (1996). Influence of dendritic structure on firing pattern in model neocortical neurons. Nature, *vol* 382, 363–366. http://www.cnl.salk.edu/CNL/

Manto, M., Bower, J. M., Conforto, A. B., Delgado-García, J. M., Da Guarda, S. N. F., Gerwig, M., Habas, C., Hagura, N., Ivry, R. B., Marien, P., Molinari, M., Naito, E., Nowak, D. A., Ben Taib, N. O., Pelisson, D., Tesche, C. D., Tilikete, C., & Timmann, D. (2012). Consensus paper: Roles of the cerebellum in motor control-the diversity of ideas on cerebellar involvement in movement. Cerebellum, 11(2), 457–487. 10.1007/s12311-011-0331-9

Markanday, A., Hong, S., Inoue, J., De Schutter, E., & Thier, P. (2023). Multidimensional cerebellar computations for flexible kinematic control of movements. Nature Communications, 14(1). 10.1038/s41467-023-37981-0

Midtgaard, J. (1992). Stellate cell inhibition of Purkinje cells in the turtle cerebellum in vitro. The Journal of Physiology, 457(1), 355–367. 10.1113/jphysiol.1992.sp019382

Mittmann, W., Koch, U., & Häusser, M. (2005). Feed-forward inhibition shapes the spike output of cerebellar Purkinje cells. Journal of Physiology, *563*(2), 369–378. 10.1113/jphysiol.2004.075028

Ohtsuki, G. (2020). Modification of synaptic-input clustering by intrinsic excitability plasticity on cerebellar purkinje cell dendrites. Journal of Neuroscience, 40(2), 267–282. 10.1523/JNEUROSCI.3211-18.2019

Ohtsuki, G., & Hansel, C. (2018). Synaptic Potential and Plasticity of an SK2 Channel Gate Regulate Spike Burst Activity in Cerebellar Purkinje Cells. IScience, 1, 49–54. 10.1016/j.isci.2018.02.001

Ohtsuki, G., Piochon, C., Adelman, J. P., & Hansel, C. (2012). SK2 channel modulation contributes to compartment-specific dendritic plasticity in cerebellar Purkinje cells. Neuron, 75(1), 108–120. 10.1016/j.neuron.2012.05.025

Ohtsuki, G., Shishikura, M., & Ozaki, A. (2020). Synergistic excitability plasticity in cerebellar functioning. In FEBS Journal (Vol. 287, Issue 21, pp. 4557–4593). Blackwell Publishing Ltd. 10.1111/febs.15355

Pouzat, C., & Hestrin, S. (1997). Developmental Regulation of Basket/Stellate Cell→Purkinje Cell Synapses in the Cerebellum. The Journal of Neuroscience, 17(23), 9104–9112. 10.1523/JNEUROSCI.17-23-09104.1997

Psarrou, M., Stefanou, S. S., Papoutsi, A., Tzilivaki, A., Cutsuridis, V., & Poirazi, P. (2014). A simulation study on the effects of dendritic morphology on layer V prefrontal pyramidal cell firing behavior. Frontiers in Cellular Neuroscience, 8. 10.3389/fncel.2014.00287

Rancillac, A., & Crépel, F. (2004). Synapses between parallel fibres and stellate cells express long-term changes in synaptic efficacy in rat cerebellum. Journal of Physiology, 554(3), 707–720. 10.1113/jphysiol.2003.055871

Rancz, E. A., & Häusser, M. (2006). Dendritic calcium spikes are tunable triggers of cannabinoid release and short-term synaptic plasticity in cerebellar purkinje neurons. Journal of Neuroscience, *26*(20), 5428–5437. 10.1523/JNEUROSCI.5284-05.2006

Rancz, E. A., & Häusser, M. (2010). Dendritic spikes mediate negative synaptic gain control in cerebellar Purkinje cells. Proceedings of the National Academy of Sciences of the United States of America, 107(51), 22284–22289. 10.1073/pnas.1008605107

Roth, A., & Häusser, M. (2001). Compartmental models of rat cerebellar Purkinje cells based on simultaneous somatic and dendritic patch-clamp recordings. Journal of Physiology, 535(2), 445–472. 10.1111/j.1469-7793.2001.00445.x

Sah, P. (1996). Ca2+-activated K+ currents in neurones: types, physiological roles and modulation. Trends in Neurosciences, *19*(4), 150–154. 10.1016/S0166-2236(96)80026-9

Shim, H. G., Jang, D. C., Lee, J., Chung, G., Lee, S., Kim, Y. G., Jeon, D. E., & Kim, S. J. (2017). Long-term depression of intrinsic excitability accompanied by synaptic depression in cerebellar purkinje cells. Journal of Neuroscience, 37(23), 5659–5669. 10.1523/JNEUROSCI.3464-16.2017

Smith, S. L., & Otis, T. S. (2005). Pattern-dependent, simultaneous plasticity differentially transforms the input-output relationship of a feedforward circuit. www.pnas.orgcgidoi10.1073pnas.0505028102

Solinas, S. M. G., Maex, R., & De Schutter, E. (2006). Dendritic amplification of inhibitory postsynaptic potentials in a model Purkinje cell. European Journal of Neuroscience, 23(5), 1207–1218. 10.1111/j.1460-9568.2005.04564.x

Steuber, V., Mittmann, W., Hoebeek, F. E., Silver, R. A., De Zeeuw, C. I., Häusser, M., & De Schutter, E. (2007). Cerebellar LTD and Pattern Recognition by Purkinje Cells. Neuron, 54(1), 121–136. 10.1016/j.neuron.2007.03.015

van Elburg, R. A. J., & van Ooyen, A. (2010). Impact of dendritic size and dendritic topology on burst firing in pyramidal cells. *PLoS Computational Biology*, 6(5), 1–19. 10.1371/journal.pcbi.1000781

van Ooyen, A., Duijnhouwer, J., Remme, M., & van Pelt, J. (2002). The effect of dendritic topology on firing patterns in model neurons. Network: Computation in Neural Systems, 13(3), 311–325. 10.1088/0954-898X/13/3/304

Vincent, P., & Marty, A. (1996). Fluctuations of inhibitory postsynaptic currents in Purkinje cells from rat cerebellar slices. The Journal of Physiology, *494*(1), 183–199. 10.1113/jphysiol.1996.sp021484

Walter, J. T., & Khodakhah, K. (2006). The linear computational algorithm of cerebellar Purkinje cells. Journal of Neuroscience, *26*(50), 12861–12872. 10.1523/JNEUROSCI.4507-05.2006

Wang, H., Dewell, R. B., Zhu, Y., & Gabbiani, F. (2018). Feedforward Inhibition Conveys Time-Varying Stimulus Information in a Collision Detection Circuit. Current Biology, 28(10), 1509–1521.e3. 10.1016/j.cub.2018.04.007

Willadt, S., Nenniger, M., & Vogt, K. E. (2013). Hippocampal feedforward inhibition focuses excitatory synaptic signals into distinct dendritic compartments. PLoS ONE, 8(11). 10.1371/journal.pone.0080984

Wulff, P., Schonewille, M., Renzi, M., Viltono, L., Sassoè-Pognetto, M., Badura, A., Gao, Z., Hoebeek, F. E., Van Dorp, S., Wisden, W., Farrant, M., & De Zeeuw, C. I. (2009). Synaptic inhibition of Purkinje cells mediates consolidation of vestibulo-cerebellar motor learning. Nature Neuroscience, 12(8), 1042–1049. 10.1038/nn.2348

Zang, Y., & De Schutter, E. (2021). The cellular electrophysiological properties underlying multiplexed coding in purkinje cells. Journal of Neuroscience, 41(9), 1850–1863. 10.1523/JNEUROSCI.1719-20.2020

Zang, Y., Dieudonné, S., & De Schutter, E. (2018). Voltage- and Branch-Specific Climbing Fiber Responses in Purkinje Cells. Cell Reports, 24(6), 1536–1549. 10.1016/j.celrep.2018.07.011

Zhou, H., Lin, Z., Voges, K., Ju, C., Gao, Z., Bosman, L. W. J., Ruigrok, T. J., Hoebeek, F. E., De Zeeuw, C. I., & Schonewille, M. (2014). Cerebellar modules operate at different frequencies. ELife, 2014(3). 10.7554/eLife.02536

