## Supplementary Material for "Purkinje cell branch morphology determines effect of inhibition and SK2 modulation on somatic pauses"

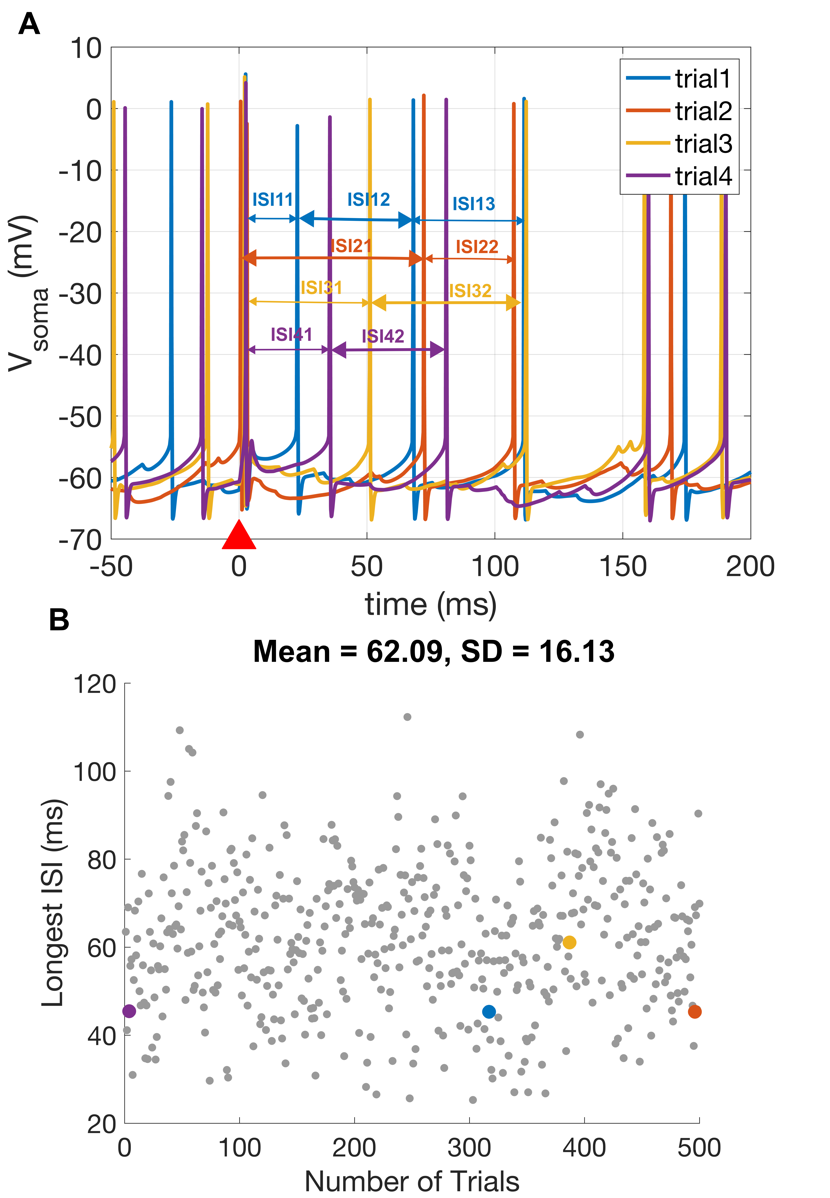

**Figure S1: Example of how ISIs are computed. Related to Figures 1, 2, 3 and 5.** Panel **A** shows a few voltage traces recorded at soma for 100PF for branch 4, for 4 different trials (blue: trial 1, red: trial 2, yellow: trial 3 and purple: trial 4). The triangle red marker (t=0) shows the time at which the PF stimulus is activated. Within a time frame of 120 ms, we compute all ISIs for all trials and we select the maximum ISI recorded in the time window examined. For example, for the first trial: ISI_trial 1 = max(ISI11,ISI12,ISI13) = ISI12. The maximum ISIs for each of the 4 trials are indicated using a thick double arrow. Panel **B** shows a scatter plot (in gray) of the maximum ISIs recorded for all 500 trials. The colored markers (blue, red, yellow and purple) represent the maximum ISIs for the four trials shown in panel A. This yields a mean of 62.09 ms, which represents the value corresponding to 100PF, which is showed in the ISI data point in Figure 1N.

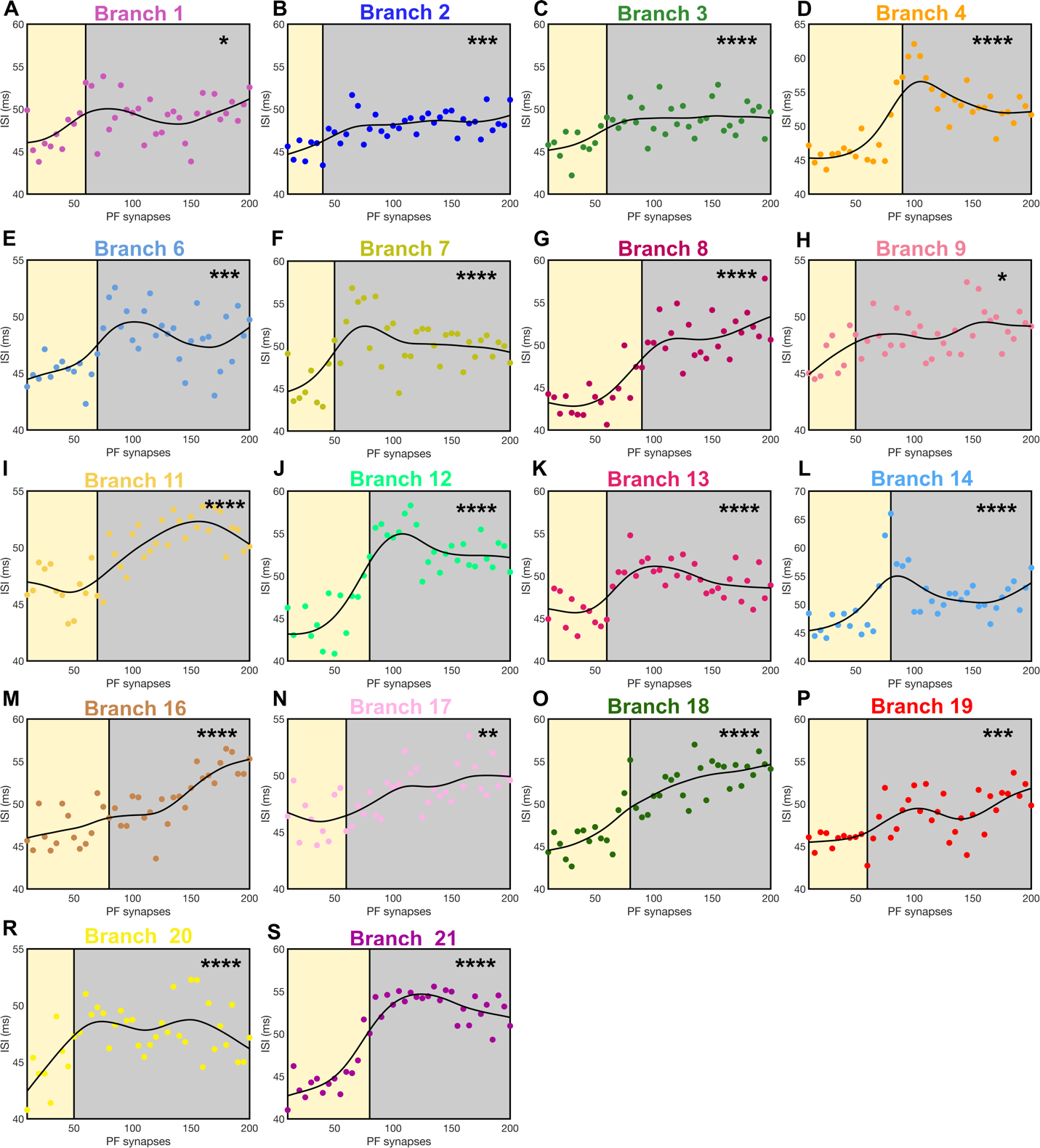

**Figure S2.** **Somatic responses: Longest ISIs for all branches. Related to Figure 2.** Scatter plots of the ISIs distribution for all branches. Each ISI data point represents the averaged values through 500 trials for each PF value. The continuous black line represents the fitted data using a smooth spline function. The yellow rectangles contain all PF numbers that are below the threshold value corresponding to each branch, while the gray rectangles comprise of all the supra-threshold PF values. The branch-specific color coding follows the same pattern shown in **Figure 1A**.

| **br #** | **hval** | **pval** | **Mean_A** | **SD_A** | **N_A** | **Mean_B** | **SD_B** | **N_B** | **Mean_diff** | **zval** | **ranksum** |
| --- | --- | --- | --- | --- | --- | --- | --- | --- | --- | --- | --- |
| 1 | 0 | 4.4e-02 | 47.611 | 2.63 | 25 | 49.249 | 2.521 | 25 | 1.638 | -2.018 | 533 |
| 2 | 1 | 4.5e-04 | 46.679 | 1.896 | 30 | 48.283 | 1.436 | 30 | 1.604 | -3.511 | 677 |
| 3 | 1 | 3.3e-05 | 46.325 | 2.098 | 25 | 48.985 | 1.891 | 25 | 2.66 | -4.152 | 423 |
| 4 | 1 | 3.5e-06 | 47.273 | 3.282 | 20 | 53.636 | 3.628 | 20 | 6.363 | -4.639 | 238 |
| 5 | 1 | 2.4e-06 | 46.748 | 3.647 | 20 | 53.136 | 1.818 | 20 | 6.388 | -4.72 | 235 |
| 6 | 1 | 6.2e-04 | 46.256 | 1.63 | 20 | 48.715 | 2.427 | 20 | 2.459 | -3.422 | 283 |
| 7 | 1 | 9.3e-05 | 46.756 | 2.573 | 20 | 51.24 | 3.186 | 20 | 4.484 | -3.909 | 265 |
| 8 | 1 | 3.0e-07 | 44.553 | 2.289 | 20 | 51.065 | 2.246 | 20 | 6.512 | -5.126 | 220 |
| 9 | 0 | 2.1e-02 | 46.799 | 1.665 | 20 | 48.294 | 1.862 | 20 | 1.495 | -2.313 | 324 |
| 10 | 0 | 2.1e-02 | 46.746 | 1.823 | 25 | 48.143 | 2.083 | 25 | 1.397 | -2.309 | 518 |
| 11 | 1 | 3.1e-06 | 46.491 | 1.915 | 20 | 50.815 | 2.18 | 20 | 4.325 | -4.666 | 237 |
| 12 | 1 | 4.6e-09 | 45.75 | 2.65 | 25 | 53.451 | 2.31 | 25 | 7.701 | -5.86 | 335 |
| 13 | 1 | 2.2e-07 | 46.23 | 1.785 | 25 | 50.013 | 1.969 | 25 | 3.783 | -5.181 | 370 |
| 14 | 1 | 7.0e-05 | 48.146 | 3.968 | 25 | 51.991 | 3.125 | 25 | 3.845 | -3.978 | 432 |
| 15 | 1 | 6.8e-08 | 47.873 | 1.833 | 20 | 66.641 | 4.458 | 20 | 18.767 | -5.396 | 210 |
| 16 | 1 | 5.1e-06 | 46.958 | 2.167 | 25 | 51.373 | 3.142 | 25 | 4.415 | -4.56 | 402 |
| 17 | 1 | 1.3e-03 | 46.795 | 2.019 | 25 | 48.875 | 2.014 | 25 | 2.08 | -3.221 | 471 |
| 18 | 1 | 1.2e-08 | 46.72 | 2.123 | 25 | 53.07 | 2.921 | 25 | 6.35 | -5.704 | 343 |
| 19 | 1 | 1.5e-04 | 46.337 | 1.643 | 25 | 49.157 | 2.523 | 25 | 2.82 | -3.784 | 442 |
| 20 | 1 | 8.6e-06 | 45.037 | 1.908 | 20 | 48.419 | 1.844 | 20 | 3.382 | -4.45 | 245 |
| 21 | 1 | 7.4e-09 | 45.765 | 2.795 | 25 | 53.352 | 1.745 | 25 | 7.587 | -5.782 | 339 |
| 22 | 0 | 4.6e-01 | 45.689 | 1.907 | 25 | 45.373 | 1.739 | 25 | -0.316 | 0.737 | 676 |

**Table S1: Statistical analysis for the ISIs observed at soma. Related to Figure 1-3.** For each branch, we show the p-value, the mean of group A and group B, its corresponding SDs, the number of samples in group A and B, the mean difference between the groups and the values corresponding to the Wilcoxon test ranksum and zval. The hval on the second column is 1 if the hypothesis that the means of the two populations are significantly different is true, and 0, if it is false. Observe (in gray) that 4 of the branches have an hval = 0. Moreover, these branches also have a p-value larger than 0.01 (99% confidence level), suggesting that the mean of group A is not significantly different than the mean of group B. For these four branches we propose various mechanisms such as increasing SK2 conductance density and adding inhibition via stellate cells, such that the difference between the two means becomes significant.

| **br #** | $\boldsymbol{\alpha}_{\boldsymbol{SK}\boldsymbol{2}}$ | **hval** | **pval** | **Mean_A** | **SD_A** | **N_A** | **Mean_B** | **SD_B** | **N_B** | **Mean_diff** | **zval** | **ranksum** |
| --- | --- | --- | --- | --- | --- | --- | --- | --- | --- | --- | --- | --- |
| 1 | 1 | 0 | 4.4e-02 | 47.611 | 2.63 | 25 | 49.249 | 2.521 | 25 | 1.638 | -2.018 | 533 |
|  | 2 | 1 | 6.9e-05 | 47.35 | 2.52 | 25 | 51.36 | 2.857 | 25 | 4.009 | -3.979 | 325 |
|  | 3 | 1 | 7.8e-07 | 47.403 | 2.575 | 25 | 53.075 | 2.841 | 25 | 5.672 | -4.941 | 284 |
| 9 | 1 | 0 | 2.1e-02 | 46.799 | 1.665 | 20 | 48.294 | 1.862 | 20 | 1.495 | -2.313 | 324 |
|  | 2 | 1 | 6.6e-05 | 46.856 | 1.687 | 20 | 49.814 | 1.893 | 20 | 2.958 | -3.989 | 262 |
|  | 3 | 1 | 1.2e-06 | 46.889 | 1.711 | 20 | 50.833 | 1.904 | 20 | 3.944 | -4.855 | 230 |
| 10 | 1 | 0 | 2.1e-02 | 46.746 | 1.823 | 25 | 48.143 | 2.083 | 25 | 1.397 | -2.309 | 518 |
|  | 2 | 1 | 7.3e-06 | 46.855 | 1.756 | 25 | 50.406 | 2.195 | 25 | 3.551 | -4.484 | 331 |
|  | 3 | 1 | 1.9e-07 | 46.997 | 1.833 | 25 | 52.142 | 2.273 | 25 | 5.145 | -5.211 | 299 |
| 22 | 1 | 0 | 4.6e-01 | 45.689 | 1.907 | 25 | 45.373 | 1.739 | 25 | -0.316 | 0.737 | 676 |
|  | 2 | 0 | 1.2e-02 | 45.869 | 2.161 | 25 | 47.588 | 1.893 | 25 | 1.720 | -2.517 | 344 |
|  | 3 | 1 | 5.6e-05 | 45.926 | 2.132 | 25 | 48.935 | 1.907 | 25 | 3.001 | -4.029 | 288 |

**Table S2: Statistical analysis for SK2 modulation. Related to Figure 3.** For each branch that showed a p-value > 0.01 and an hval = 0 in **Table S1**, we show the statistical values when modulating the conductance density of the SK2 channel controlled by the parameter $\alpha_{SK2}$. As in the previous table, we show the p-value, the mean of group A and group B, its corresponding SDs, the number of samples in group A and B, the mean difference between the groups and the values corresponding to the Wilcoxon test ranksum and zval. Observe that for all branches, increasing SK2 leads to a significant increase in Mean_diff, a p-value < 0.01 and an hval = 1, denoting that all branches exhibit a somatic pause.

| **Br #** | **Stel** | **hval** | **pval** | **Mean_A** | **SD_A** | **N_A** | **Mean_B** | **SD_B** | **N_B** | **Mean_diff** | **zval** | **ranksum** |
| --- | --- | --- | --- | --- | --- | --- | --- | --- | --- | --- | --- | --- |
| 1 | 0 | 0 | 4.4e-02 | 47.611 | 2.63 | 25 | 49.249 | 2.521 | 25 | 1.638 | -2.018 | 533 |
| 1 | 75 | 1 | 1.1e-03 | 48.042 | 2.59 | 25 | 50.85 | 2.775 | 25 | 2.808 | -3.26 | 469 |
| 1 | 150 | 1 | 1.4e-04 | 48.96 | 2.576 | 25 | 52.61 | 3.084 | 25 | 3.65 | -3.803 | 441 |
| 2 | 0 | 1 | 2.3e-04 | 46.679 | 1.896 | 30 | 48.424 | 1.506 | 30 | 1.745 | -3.689 | 665 |
| 2 | 75 | 1 | 6.5e-09 | 47.158 | 1.88 | 30 | 50.985 | 1.635 | 30 | 3.827 | -5.803 | 522 |
| 2 | 150 | 1 | 3.2e-09 | 48.277 | 1.92 | 30 | 52.529 | 1.949 | 30 | 4.252 | -5.921 | 514 |
| 3 | 0 | 1 | 5.4e-05 | 46.325 | 2.098 | 25 | 48.996 | 1.96 | 25 | 2.67 | -4.036 | 429 |
| 3 | 75 | 1 | 2.5e-07 | 46.822 | 2.067 | 25 | 50.611 | 1.858 | 25 | 3.79 | -5.161 | 371 |
| 3 | 150 | 1 | 2.6e-08 | 47.652 | 2.106 | 25 | 52.274 | 2.095 | 25 | 4.622 | -5.569 | 350 |
| 4 | 0 | 1 | 5.9e-06 | 47.273 | 3.282 | 20 | 51.961 | 2.038 | 20 | 4.688 | -4.531 | 242 |
| 4 | 75 | 1 | 6.0e-07 | 47.461 | 2.338 | 20 | 53.615 | 2.268 | 20 | 6.154 | -4.991 | 225 |
| 4 | 150 | 1 | 2.2e-07 | 48.446 | 2.324 | 20 | 55.252 | 2.186 | 20 | 6.806 | -5.18 | 218 |
| 5 | 0 | 1 | 4.5e-06 | 46.748 | 3.647 | 20 | 52.895 | 1.873 | 20 | 6.146 | -4.585 | 240 |
| 5 | 75 | 1 | 3.4e-07 | 47.593 | 3.212 | 20 | 54.54 | 2.077 | 20 | 6.947 | -5.099 | 221 |
| 5 | 150 | 1 | 3.0e-07 | 48.658 | 3.155 | 20 | 55.916 | 2.298 | 20 | 7.258 | -5.126 | 220 |
| 6 | 0 | 1 | 2.3e-03 | 46.256 | 1.63 | 20 | 48.438 | 2.468 | 20 | 2.182 | -3.043 | 297 |
| 6 | 75 | 1 | 3.4e-04 | 46.979 | 1.73 | 20 | 50.085 | 2.472 | 20 | 3.107 | -3.584 | 277 |
| 6 | 150 | 1 | 8.4e-04 | 48.081 | 1.92 | 20 | 51.106 | 2.468 | 20 | 3.025 | -3.341 | 286 |
| 7 | 0 | 1 | 1.6e-04 | 46.756 | 2.573 | 20 | 51.114 | 3.228 | 20 | 4.358 | -3.773 | 270 |
| 7 | 75 | 1 | 3.7e-05 | 47.557 | 2.635 | 20 | 52.656 | 3.176 | 20 | 5.098 | -4.125 | 257 |
| 7 | 150 | 1 | 4.7e-05 | 48.802 | 2.731 | 20 | 53.885 | 3.292 | 20 | 5.084 | -4.071 | 259 |
| 8 | 0 | 1 | 3.0e-07 | 44.553 | 2.289 | 20 | 51.451 | 2.692 | 20 | 6.898 | -5.126 | 220 |
| 8 | 75 | 1 | 1.4e-07 | 45.614 | 1.993 | 20 | 53.083 | 2.542 | 20 | 7.468 | -5.261 | 215 |
| 8 | 150 | 1 | 2.6e-07 | 47.204 | 2.015 | 20 | 54.236 | 2.771 | 20 | 7.032 | -5.153 | 219 |
| 9 | 0 | 0 | 2.1e-02 | 46.799 | 1.665 | 20 | 48.294 | 1.862 | 20 | 1.495 | -2.313 | 324 |
| 9 | 75 | 1 | 3.6e-03 | 47.467 | 1.76 | 20 | 49.449 | 1.805 | 20 | 1.982 | -2.908 | 302 |
| 9 | 150 | 1 | 3.6e-03 | 48.59 | 1.747 | 20 | 50.546 | 1.922 | 20 | 1.956 | -2.908 | 302 |
| 10 | 0 | 0 | 1.1e-02 | 46.746 | 1.823 | 25 | 48.193 | 1.915 | 25 | 1.447 | -2.542 | 506 |
| 10 | 75 | 1 | 1.7e-04 | 47.39 | 1.811 | 25 | 49.763 | 2.044 | 25 | 2.373 | -3.764 | 443 |
| 10 | 150 | 1 | 3.6e-05 | 48.14 | 1.984 | 25 | 51.122 | 2.051 | 25 | 2.983 | -4.133 | 424 |
| 11 | 0 | 1 | 1.7e-07 | 46.491 | 1.915 | 20 | 51.393 | 1.678 | 20 | 4.902 | -5.234 | 216 |
| 11 | 75 | 1 | 2.2e-07 | 47.221 | 1.913 | 20 | 52.858 | 1.657 | 20 | 5.637 | -5.18 | 218 |
| 11 | 150 | 1 | 1.9e-07 | 48.306 | 2.031 | 20 | 54.109 | 1.712 | 20 | 5.803 | -5.207 | 217 |
| 12 | 0 | 1 | 6.6e-09 | 45.75 | 2.65 | 25 | 52.81 | 2.304 | 25 | 7.06 | -5.801 | 338 |
| 12 | 75 | 1 | 1.4e-09 | 46.496 | 2.356 | 25 | 54.652 | 2.32 | 25 | 8.156 | -6.054 | 325 |
| 12 | 150 | 1 | 1.6e-09 | 48.317 | 2.698 | 25 | 56.423 | 2.237 | 25 | 8.106 | -6.034 | 326 |
| 13 | 0 | 1 | 2.2e-07 | 46.23 | 1.785 | 25 | 50.013 | 1.969 | 25 | 3.783 | -5.181 | 370 |
| 13 | 75 | 1 | 1.5e-07 | 47.282 | 1.93 | 25 | 51.646 | 2.049 | 25 | 4.363 | -5.258 | 366 |
| 13 | 150 | 1 | 1.0e-06 | 48.911 | 2.183 | 25 | 53.273 | 2.367 | 25 | 4.361 | -4.89 | 385 |
| 14 | 0 | 1 | 7.0e-05 | 48.146 | 3.968 | 25 | 51.991 | 3.125 | 25 | 3.845 | -3.978 | 432 |
| 14 | 75 | 1 | 5.6e-06 | 48.548 | 2.938 | 25 | 52.973 | 3.132 | 25 | 4.426 | -4.54 | 403 |
| 14 | 150 | 1 | 8.2e-05 | 49.74 | 2.573 | 25 | 53.397 | 3.116 | 25 | 3.657 | -3.939 | 434 |
| 15 | 0 | 1 | 6.8e-08 | 47.873 | 1.833 | 20 | 66.651 | 4.253 | 20 | 18.778 | -5.396 | 210 |
| 15 | 75 | 1 | 6.8e-08 | 48.405 | 1.837 | 20 | 67.485 | 4.324 | 20 | 19.08 | -5.396 | 210 |
| 15 | 150 | 1 | 6.8e-08 | 49.633 | 1.916 | 20 | 67.647 | 4.611 | 20 | 18.014 | -5.396 | 210 |
| 16 | 0 | 1 | 5.1e-06 | 46.958 | 2.167 | 25 | 51.373 | 3.142 | 25 | 4.415 | -4.56 | 402 |
| 16 | 75 | 1 | 2.2e-06 | 47.706 | 2.182 | 25 | 52.151 | 3.059 | 25 | 4.445 | -4.734 | 393 |
| 16 | 150 | 1 | 4.3e-06 | 48.89 | 2.28 | 25 | 53.263 | 3.137 | 25 | 4.373 | -4.598 | 400 |
| 17 | 0 | 1 | 2.5e-04 | 46.795 | 2.019 | 25 | 49.124 | 1.866 | 25 | 2.329 | -3.667 | 448 |
| 17 | 75 | 1 | 1.5e-04 | 47.619 | 1.976 | 25 | 50.216 | 1.836 | 25 | 2.597 | -3.784 | 442 |
| 17 | 150 | 1 | 5.1e-04 | 48.605 | 2.223 | 25 | 51.02 | 1.743 | 25 | 2.416 | -3.473 | 458 |
| 18 | 0 | 1 | 2.6e-09 | 46.72 | 2.123 | 25 | 54.49 | 3.015 | 25 | 7.771 | -5.957 | 330 |
| 18 | 75 | 1 | 1.6e-09 | 47.399 | 2.031 | 25 | 55.031 | 2.186 | 25 | 7.632 | -6.034 | 326 |
| 18 | 150 | 1 | 2.0e-09 | 48.614 | 2.198 | 25 | 56.062 | 2.254 | 25 | 7.448 | -5.995 | 328 |
| 19 | 0 | 1 | 3.6e-05 | 46.337 | 1.643 | 25 | 49.512 | 2.519 | 25 | 3.174 | -4.133 | 424 |
| 19 | 75 | 1 | 2.9e-06 | 47.08 | 1.628 | 25 | 50.829 | 2.644 | 25 | 3.749 | -4.676 | 396 |
| 19 | 150 | 1 | 3.5e-06 | 48.202 | 1.614 | 25 | 52.102 | 2.683 | 25 | 3.899 | -4.637 | 398 |
| 20 | 0 | 1 | 3.7e-05 | 45.037 | 1.908 | 20 | 48.327 | 2.136 | 20 | 3.29 | -4.125 | 257 |
| 20 | 75 | 1 | 1.6e-06 | 45.82 | 1.924 | 20 | 50.295 | 2.194 | 20 | 4.475 | -4.801 | 232 |
| 20 | 150 | 1 | 1.4e-06 | 47.013 | 1.931 | 20 | 51.694 | 2.321 | 20 | 4.681 | -4.828 | 231 |
| 21 | 0 | 1 | 7.4e-09 | 45.765 | 2.795 | 25 | 53.479 | 1.764 | 25 | 7.714 | -5.782 | 339 |
| 21 | 75 | 1 | 1.4e-09 | 46.753 | 1.781 | 25 | 55.049 | 1.862 | 25 | 8.296 | -6.054 | 325 |
| 21 | 150 | 1 | 3.7e-09 | 48.748 | 2.29 | 25 | 56.448 | 1.952 | 25 | 7.701 | -5.898 | 333 |
| 22 | 0 | 0 | 3.7e-01 | 45.689 | 1.907 | 25 | 45.321 | 1.681 | 25 | -0.368 | 0.893 | 684 |
| 22 | 75 | 0 | 5.5e-01 | 47.661 | 2.291 | 25 | 47.225 | 1.76 | 25 | -0.436 | 0.601 | 669 |
| 22 | 150 | 0 | 1.2e-02 | 50.205 | 2.528 | 25 | 48.604 | 1.864 | 25 | -1.6 | 2.503 | 767 |

**Table S3:** **Statistical analysis of ISI when considering feed-forward inhibition via stellate cells. Related to Figure 5.** For each branch, we show the p-value, the mean of group A and group B, its corresponding SDs, the number of samples in group A and B, the mean difference between the groups and the values corresponding to the Wilcoxon test ranksum and zval. In gray shade we show the branches for which group A and group B are not significantly different (hval =0) as shown by the p-value larger than 0.01. Observe that for branch 1, 9 and 10, adding inhibition results in a meaningful statistical difference between the two populations. Only exception is branch 22: for maximum studied inhibition (150 stellate cell synapses), results in a p-value of 0.012, which is still rejected by the 99% confidence level used in our statistics.
